# Determination of factors that allow cryogenic nanoscopy with high power illumination without devitrification

**DOI:** 10.1101/2025.09.26.678528

**Authors:** Jan Huebinger, Philippe I.H. Bastiaens

**Author notes:** Author deceased (Philippe I.H. Bastiaens).

## Abstract

Cryogenic super-resolution fluorescence light microscopy, or nanoscopy, has been demonstrated to be useful to close the resolution gap in cryogenic correlative light and electron microscopy (CLEM). Importantly, under cryogenic conditions fundamental resolution barriers that are imposed by molecular motion and photobleaching on fluorescence microscopy are circumvented. The resolution gain is even higher in nanoscopic methods, due to higher obtainable resolution and slower acquisition speed, rendering cryo-nanoscopy promising even beyond CLEM. However, cryogenic nanoscopy is often limited by heating and devitrification of the sample through strong laser irradiation, with drastic differences in reported tolerable power densities. We therefore investigated the laser-induced heating in different setups for cryogenic nanoscopy by time-dependent finite-element simulations complemented with absorption measurements of mammalian cells. Laser-induced heating happened in milliseconds, precluding efficient sample preservation by most intermittent illuminations. Under moderate light densities used for single molecule localization microscopy, absorbance by mammalian cells was too weak to explain devitrification and heating is governed by absorption of supporting material. However, the much higher power densities used in stimulated emission depletion nanoscopy resulted in temperatures clearly above the devitrification temperature from absorption by cells alone, unless the sample was mounted on an efficient heat exchanger, such as a diamond.

---

While cryo-fluorescence microscopy has been proven useful for cryogenic correlative light and electron microscopy/tomography (CLEM) ^1-4^, it has been also shown that cryogenic fluorescence microscopy breaks through a fundamental resolution barrier imposed by motional blur and photobleaching ^5^, rendering it also a very auspicious method for fluorescence micro- and nanoscopy outside of CLEM.

In order to cryo-arrest living cells in a ‘close-to-native’ state, they have to be cooled extremely rapidly below the glass transition temperature of water to avoid destructive ice crystal formation in a process called vitrification ^6,7^. They further have to be kept below the glass transition temperature during the whole preparation and imaging process to avoid devitrification. The absorption of energy from the irradiation is an inevitable source of heating during fluorescence micro- or nanoscopy, which can be strong enough to lead to ice crystallization ^3^.

Upon heating above the glass transition temperature of water, an aqueous sample will first become fluid and then crystallize, if it remains sufficiently long between the glass transition and the melting temperature ^8^. At temperatures close to the glass transition temperature, ice crystallization is relatively slow ^9^ and vitrification of aqueous solutions heated to this temperature regime can reoccur from a liquid state upon re-cooling ^10^. Revitrification can also occur upon re-cooling, when a sample is directly heated above the melting temperature by laser irradiation ^9^. However, any heating of the sample above the glass transition temperature of water leads to a drastic increase in molecular motility and thus to changes in the sample, such as diffusion of molecules or conformational changes in proteins ^11^. Since localization measurements with sub-nanometer precision and measurements of protein conformations are nowadays feasible by cryogenic fluorescence microscopy ^5,12^ and cryo-electron tomography ^13^, such changes should be avoided.

Single molecule localization microscopy (SMLM) ^14^ has been used to close the resolution gap between fluorescence light microscopy and electron microscopy (EM) ^3,15^. Since photobleaching has been shown to be dramatically reduced under cryogenic conditions ^2,5,12^, resolution in SMLM could be markedly improved under cryogenic conditions, despite the use of air objectives with relatively low numerical aperture (NA) ^12^. However, it has also been noticed that prolonged exposure to laser light during typical SMLM illumination schemes led to heating above the glass transition temperature of water, as evident from ice crystallization in the sample ^3^. Ice crystallization was thereupon circumvented by lower illumination powers, intermittent irradiation profiles and/or addition of cryoprotective agents to the sample ^3,16-19^. However, the use of cryoprotective agents is prone to change the state of living cells even before they are cryo-fixed ^10,20^ and reducing illumination powers limits the applicability of several nanoscopy methods. Further, the absence of ice crystals is not sufficient to prove that the sample is kept below the glass transition temperature of water. Changes to the sample, such as diffusion or protein conformational changes, can thus not be excluded by the absence of ice crystals. Recently, the thermal properties of EM grids under illumination have been studied in more detail by a combination of experiment and steady-state heat transfer simulations. By this, alternative support material was found that led to significantly less heating during SMLM ^21-23^.

More recently, stimulated emission depletion (STED) nanoscopy ^24^ has been used under cryogenic conditions. These conditions enabled nanoscopy measurements in cells that were not possible at room temperature due to inhibition of motional blur and drastic reduction of photobleaching ^5^. STED nanoscopy has the advantage of being directly (semi-)quantitative and is thus of great interest for quantitative fluorescence microscopy. However, it comes at the cost of much higher irradiation densities. Despite irradiation density (∼10^7^ Wcm^-2^) that were approximately 5 orders of magnitude higher than those used for cryo-SMLM, no change in the sample was observed during continuous STED nanoscopy acquisition ^5^. However, the state of the water was not directly measured, since no subsequent EM was performed.

One explanation for this apparent discrepancy in sample heating between the SMLM and STED nanoscopy measurements could be different sample mounting. For cryo-STED nanoscopy a thin (10-15 μm) aqueous sample containing living cells was mounted directly on a diamond heat exchanger of superior thermal conductivity (>10000 Wm^-1^K^-1^ at −196°C)^25^. Cryo-SMLM was performed on an EM grid placed in vacuum or gar environment, where the heat had to be conducted laterally a relatively long distance through the thin film of aqueous sample of low thermal conductivity (<1 Wm^-1^K^-1^) or the support film material. On the other hand, the strongly focused (∼1 μm^2^) STED beam has a much greater surface to volume ratio then the widefield illumination used in SMLM (e.g. 100 μm^2^ in ^3^), which can also contribute to faster heat dissipation. Additionally, illumination for SMLM is static and lasts for a relatively long time (typically several minutes), whereas the STED beam is scanned over the sample, so that each point in the sample is typically only directly illuminated for few to tens of milliseconds.

We therefore sought to unravel, which factors govern heating and devitrification upon laser irradiation. Additionally, we sought to investigate, if there is heating above the glass transition temperature in those conditions, where ice crystallization was not detectable or not tested. Lastly, we wanted to estimate the heating induced by STED nanoscopy on EM grids, to estimate if STED nanoscopy without heating above the glass transition temperature is feasible in this configuration.

To achieve this, we performed time-dependent finite-element simulations of laser-light absorption in the different sample configurations coupled to conductive heat transfer within those samples. To assess direct heating of mammalian cells by the different laser irradiations, we also determined their absorption coefficient experimentally.

Time-dependent simulations showed that steady-state temperatures are generally reached in tens of millisecond after onset of irradiation. Thus, previously employed intermittent irradiation profiles still led to heating above the glass transition temperature. We measured the absorption coefficient of mammalian cells to be 4 and 2 orders of magnitude higher than water at visible and near infrared irradiation wavelengths, respectively. Simulations of SMLM on EM grids showed that heating is nevertheless dominated by absorption of the support film and absorbance by mammalian cells is not sufficient for heating above the glass transition temperature. Our simulations confirm the reduced heating on holey gold or silver-coated carbon film ^21-23^. This is however mostly achieved by the high reflection of these films, which can be limiting for fluorescence micro- and nanoscopy. We therefore tested further film material and found SiO_2_ films as very promising low absorption material that would enable >100-fold stronger illumination. However, even without illumination of the support film, intense STED illumination would heat cells on EM grids in an area of tens of micrometer around the scanning beam above the glass transition temperature. This heating is drastically mitigated, when cells are mounted on a diamond heat exchanger 5, where temperatures stayed clearly below the glass transition temperature.

## Results

### Heating of aqueous samples by SMLM and STED nanoscopy

We simulated first the heating in the central point of a 0.5-μm aqueous sample on an EM grid with a holey carbon support film during SMLM illumination in a sample geometry as previously described ^3,26^(see method, figure 1a). To approximate an aqueous sample, we used published absorption values of water, the main constituent of buffers and cells. Upon laser irradiation with 0.3 mW at 488-nm plus 0.015 mW of 405-nm over an area of 100 μm^2^ (=300 + 15 Wcm^-2^), a new steady-state temperature of −100 °C with an expected temperature gradient towards the periphery emerged. The steady-state temperature is approached by <1 °C difference within <20 ms (Figure 1 b,c; Supplementary Movie 1). This temperature is clearly above the glass transition temperature of water, where ice crystals will form, yet crystal growth is still relatively slow ^9,10^. However, when the supporting carbon film was omitted in the simulation, temperature increased by less than 1 °C and thus stayed far below the glass transition temperature of water. We therefore also investigated the effects of alternative support films. The higher transparency of formvar and silicon monoxide films had been shown to allow a higher irradiation density, but are also considered suboptimal for fluorescence microscopy and subsequent EM ^16,18,19,22^. Commercially available holey 50-nm gold support films ^27^ and custom 50-nm silver-coating on holey carbon films ^21^ have been shown to heat less than holey carbon films ^21-23^. We confirmed that both of the films heat much less than holey carbon (Figure 1c). It should however be noted that the reduction in heating comes mainly from the strong reflection of gold (44%) and silver (95%). The gold and silver films actually absorb >90% and >97% of the light that is not reflected on their surface, resulting in very low transmission of these films (Figure 1d). This will mask information on parts of the sample that is not in the holes and strong reflections would impose strong requirements on the filtering of the detected light.

**Figure 1:**
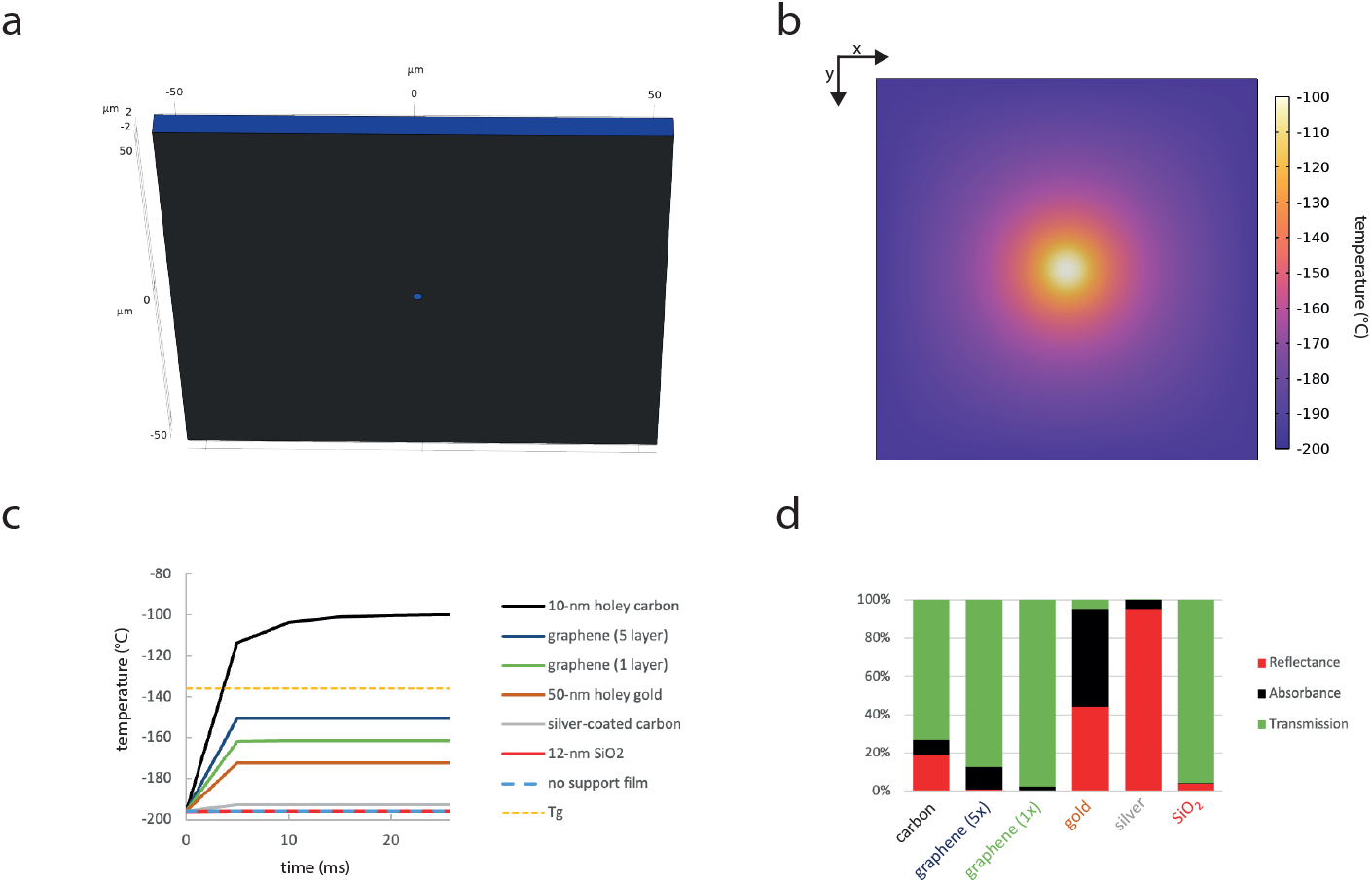
Heating of aqueous samples on EM grids by SMLM. a) The sample represents an individual square of an EM grid, where a 0.5-5 μm aqueous or cell layer (blue) was placed on a carbon film (black) with a 2-μm hole in the center. b) x-y view of the temperature profile of the EM grid close to steady-state after 40 ms of SMLM laser irradiation (0.315 mW) at a 100-μm^2^ spot in the center. c) Left: Temperature over time in the central point of the water layer from start of the SMLM laser irradiation in the EM grid configuration using a 10-nm amorphous carbon support (black line), a 50-nm gold support (red line), a 12-nm SiO_2_ support (red line), 1 or 5 layers of graphene (dark blue and green lines), a 50-nm silver-coating on an amorphous carbon support (grey) or no support film (dashed blue line); Tg: glass transition temperature of water (dashed yellow line). Right: Corresponding transmissions of the support films at 488-nm

We therefore also tested SiO2 films ^28^ and continuous graphene films ^29^, which have much better transmissivity (Figure 1d). Graphene has, despite its atomically thin layer (0.34 nm), a relatively high absorption of 2.3% per atomic layer ^30^, but can have very high in-plane thermal conductivity of >2000 Wm^-1^K^-1 31^. Graphene is typically used on top of a holey carbon film ^29,32^, which would not change the heating properties a lot, but is also commercially available directly on EM grids. Simulations showed that despite a minimal reflectivity ^33^, aqueous samples on 1 or 5 layers of graphene were not heated above the glass transition temperatures. Here, it has to be however noted, that exact thermal conductivities of modified graphene layers that are typically used ^27,32^ are not exactly known (see Methods section). SiO_2_ support films can be around 12 nm thin ^28^ and have an orders of magnitude lower absorption coefficient (4.5×10^4^ m^-1^) and much lower reflection (∼4%) at 488-nm compared to the other support film materials. A 12-nm amorphous SiO_2_ support film led to less than 1°C stronger heating compared to a sample without support film (Figure 1c) and laser intensities could be increased by a factor of 100 before the glass transition temperature of water was reached (Supplementary figure 1).

We next simulated the heating of a STED laser focused into an aqueous sample mounted to a diamond heat exchanger on the cryo-arrest stage. For this, the sample geometry was constructed as described ^5^(see methods, figure 2a). Here, a constant irradiation with 1 W of 775-nm laser light at the central 1.5-μm^2^ spot (∼6.7×10^7^ Wcm^-2^) did not raise the temperature by more than 1 °C in this spot (Figure 2 b,c).

**Figure 2:**
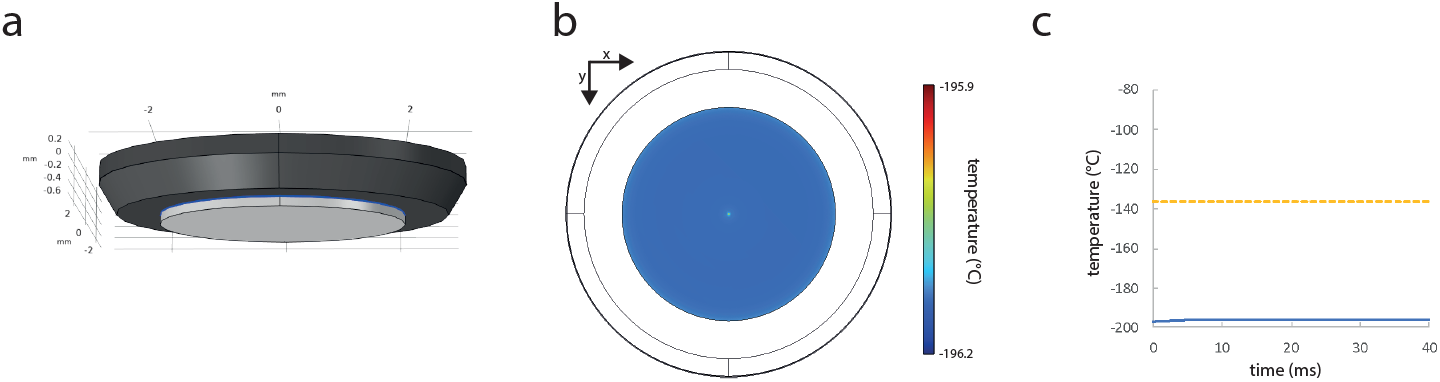
Heating of aqueous samples on a diamond heat exchanger by STED nanoscopy. a) The cryo-arrest sample was built as a 10-μm aqueous or cell layer (blue) between a cover slide (light grey) and a diamond (dark grey). b) x-y view of the temperature profile of the water layer close to steady-state after 40 ms of STED laser irradiation (1 W) at a 1.5-μm^2^ spot in the center. c) Temperature over time in the central point of the water layer from start of the STED laser irradiation (blue line); glass transition temperature of water is indicated by the dashed yellow line.

### Measurement of absorption coefficient of mammalian cells

Cells absorb significantly more visible light than water, as evident from absorption measurements of cell suspensions ^34,35^, and will therefore be heated more strongly than water by laser irradiation. However, the simulation of heating of cells upon laser irradiation requires the linear absorption coefficient of the cells. This absorption coefficient cannot be extrapolated from suspension measurements, because the linear relationship between concentration and absorption is only valid for solutions and not suspensions ^36-38^. We therefore sought to measure the mean absorption coefficient of mammalian cells experimentally.

To this end we cultured MDCK cells in confined monolayers and measured the extinction coefficient in a transmission light microscope (Figure 3a) together with measurements of cell heights by confocal laser scanning microscopy on the same samples (Supplementary figure 2). Using a 20x 0.75NA objective, a coefficient of 639 ± 211 m^-1^ (mean ± sd; n=6) was determined. However, this extinction coefficient contains not only absorption, but also part of the scattering that missed the objective. It can therefore only demark an upper limit to the absorption coefficient. Scattering of mammalian A375 cells and lymphocytes has been measured in dependence of the scattering angle ^39^ (Supplementary figure 3). We therefore designed an experiment, where we used non-convergent illumination and measured the extinction coefficient with two different objectives with different NAs, i.e. two different collection angles. Using this approach and the published angle-dependent scattering, we were able to disentangle the scattering from the absorption coefficient (figure 3b; Methods section), resulting in an absorption coefficient of 289 ± 114 m^−1^(mean ± sem; N=7; n=68-162), which is 4 orders of magnitude higher than the absorption of water at 488-nm.

**Figure 3:**
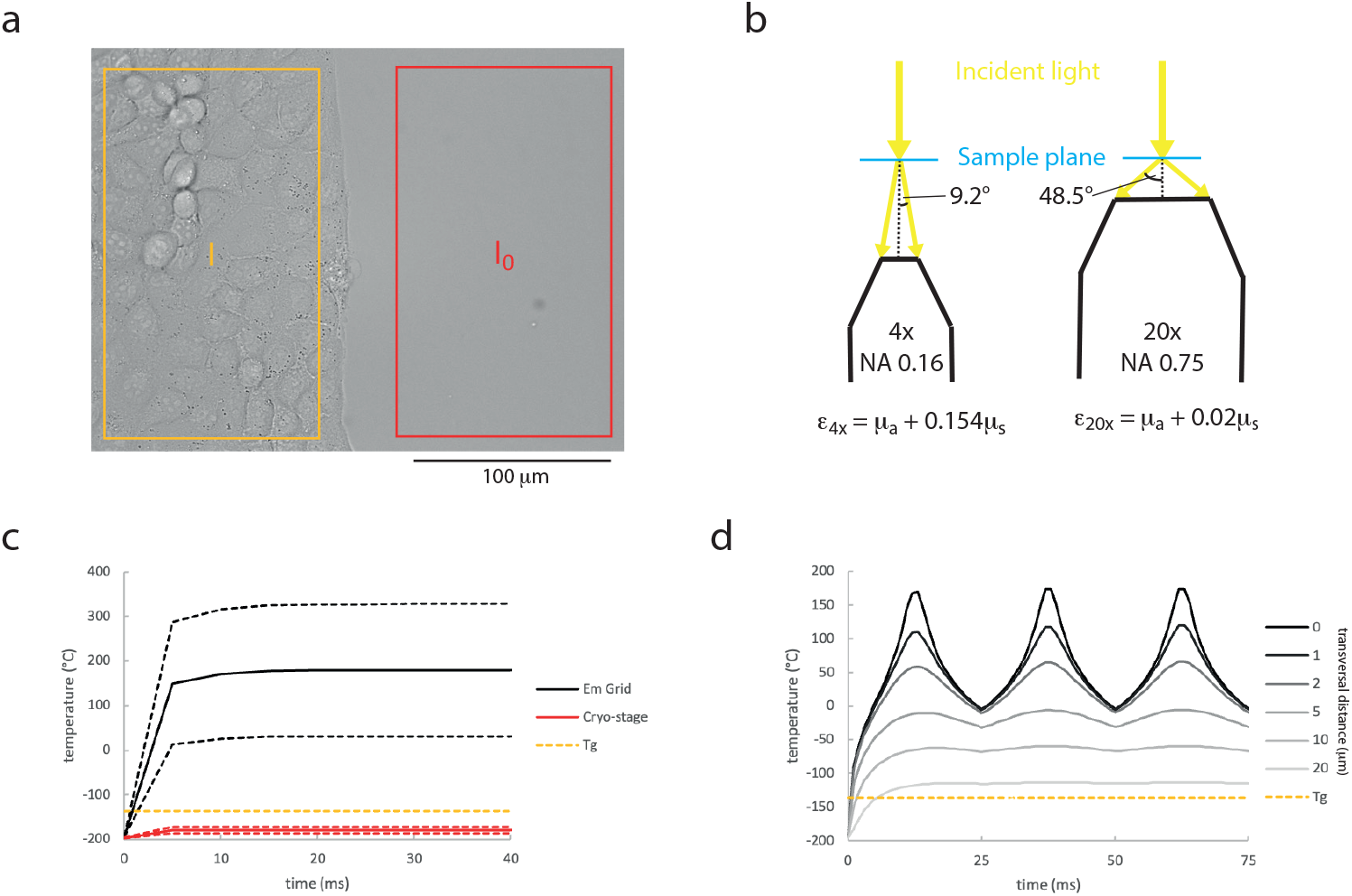
Absorption and heating of mammalian cells. a) Representative image of an extinction coefficient measurement using a 20x 0.75 NA objective at the border of a confined monolayer of MDCK cells. The transmitted light intensity after extinction by the cell monolayer (I) and the reference light intensity (I_0_) are measured in the same field of view. b) Schematic of the experiment for measuring the absorption coefficient of cells. Extinction coefficients (*ε*) were measured with a 4x 0.16 NA (left) and a 20x 0.75 NA (right) under non-convergent illumination. The extinction coefficient of the cells is composed of the absorption coefficient (*μ*_*a*_) and the fraction of the scattering coefficient (*μ*_*s*_) that misses the objective, i.e. that has a scattering angle larger than 9.2° (4x; 15.4±0.3%) or 48.5° (20x; 2±1%). c) Temperature over time in the central point of a cell layer (linear absorption coefficient 289±114 m^-1^) from start of laser irradiation by a static 1-W STED laser focused to a 1.5-μm^2^ spot (∼6.7×10^7^ W/cm^2^) in the EM grid configuration omitting a support film (black lines) and on the ultra-rapid cryo-arrest stage (red lines); Tg: glass transition temperature of water (yellow dashed line) d) Evolution of temperature of a cellular sample (absorption coefficient: 289 m^-1^) on an EM grid during 3 lines of scanning (length: 10-μm) with 40 nm transversal spacing by a 1-W STED laser. The temperature is depicted at the center of the 2^nd^ scanning line (0) and at indicated transversal distances to the measurement point (gray lines). Tg: glass transition temperature of water (yellow dashed line)

Using this absorption coefficient, we first simulated the heating of the cell by SMLM. When no support film was considered, this resulted in a heating of less than 2°C (Supplementary figure 4).

We then simulated heating by a static STED laser of a cell mounted to the diamond heat exchanger of the cryo-stage versus a cell on an EM grid without any support film. The latter would be a realistic scenario, if the scanning was confined to a hole in a holey support film. The temperature in the cellular sample on the cryo-stage stayed clearly below the glass transition temperature of water. However, on an EM grid, temperatures were reached that would melt or even evaporate the sample locally (figure 3c). It should be noted that above the melting temperature, convective heat transport becomes a significant factor. Therefore, the temperature development above 0°C cannot be reliably simulated by a conductive heat-transfer model. Further, in a real STED experiment, the laser is not static, but constantly scanning over the sample. We therefore also simulated scanning of the laser over a 10 μm wide field of view of the sample on an EM grid with a pixel length of 40 nm and a pixel dwell time of 100 μs (Supplementary Movie 2). Under these conditions temperatures approached the steady-state temperatures within a few °C when the scanning beam directly hits a region of the sample and the surrounding sample is constantly heated above the glass transition temperature within a distance >20 ± 10 μm (figure 3d, Supplementary figure 5).

## Discussion

To our knowledge, there is no previous report on the linear absorption coefficient of mammalian cells. We have therefore determined this value experimentally. Cellular absorption thereby tends to be higher than tissue absorption (3-160 m^-1^) ^40^, which could be explained by the interstitial spaces absorbing less than the cells themselves. Also, a 30-40x higher scattering than absorption coefficient is consistent between cells and tissues. The relatively large uncertainty for the obtained cellular absorption originates in large parts from the large relative differences in angle dependent scattering (∼ ±50% at 50°) between different cell types ^39^, which propagates as uncertainty into the absorption coefficient. The result therefore also reflects differences between cell types. Furthermore, it is to be expected that there are local absorption differences over individual cells due to sub-cellular spatial inhomogeneity. Despite these uncertainties, the results give clear indications under which conditions heating above the glass transition temperatures will occur.

Chang et al. found ice formation on EM grids with carbon support films only after prolonged exposure time ^3^. This is in very good agreement with the rapidly established steady-state temperature of −100°C. At such low temperatures, ice formation is slow ^10^. It is therefore consistent that ice formation could be avoided by the use of cryoprotective agents and intermittent illumination schemes ^3^. It is also in good agreement with other published results, where the threshold for long exposure times without ice formation in the absence of cryoprotective agents was found between 30 and 100 Wcm^-2 17-19^, which would correspond to a laser power of 0.03 – 0.1 mW in the tested setup. The absolute tolerable intensity will thereby also depend on excitation wavelength and illumination area. However, these factors appear to be relatively minor and all results are within the same order of magnitude.

Clearly, ice crystallization needs to be avoided in CLEM, since it is detrimental for EM. Furthermore, ice-crystals are also detrimental for fluorescence micro- or nanoscopy outside of CLEM, since water concentrates in ice-crystals and displaces other molecules, leading to distortion of the sample ^41^. Importantly, when samples are heated above the glass transition temperature of water without sufficient time to crystallize, diffusional and conformational changes in the sample are still occurring. This will also alter the sample in an uncontrolled way. Given that high-resolution data is nowadays obtainable by cryo-electron tomography, cryo-fluorescence nanoscopy or cryo-microspectroscopy ^5,12,13^, such changes can readily translate into distortions of the results. Our results show that temperature steady-states upon laser irradiation are approached within <10 ms. This shows that intermittent irradiation profiles will not prevent the sample from reaching temperatures above the glass transition, but they only prohibit ice crystallization by reducing the time segments above the glass transition. It is of note that the success of avoiding ice crystallization by intermittent illumination ^3,16^ therefore implies that it is not only integrated time above the glass transition temperature that governs ice crystallization, but the length of individual continuous time segments above the glass transition temperature appears to have a significant influence.

The presented results highlight the need of considering sample geometries for cryogenic fluorescence light microscopy. The use of highly absorbing support films should be avoided to prevent heating during fluorescence microscopy on EM grids ^16,21-23^. However, the choice of support film material for CLEM is a multifactored problem, since support films are ideally very stable under the electron beam, ensure good sample adhesion, show low autofluorescence and high transmissivity ^18,19,22,42^. Using the appropriate support film, allows a broad range of SMLM techniques to be used. In this context, it is of interest that a SiO^2^ support film allows a >100x higher illumination compared to standard holey carbon film, while maintaining a much higher transmissivity than silver or gold films.

However, for the use of strongly focused high intensity laser light, as used in STED nanoscopy, the additional mounting of the sample on an efficient heat exchanger appears necessary. Apart from the tested configuration where the sample is mounted to a heat exchanger made of diamond, which is cooled from the side opposing the sample directly by a flow of liquid nitrogen, also the mounting of a sample on a sapphire disc ^4^ can be considered. However, at the temperature of liquid nitrogen, diamond is the superior thermal conductor with thermal conductivities >10000 Wm^-1^K^-1 25^ compared to sapphire (<1000 Wm^-1^K^-1^), which reaches its peak conductivity between 10 and 30 K ^43^ and can be a very efficient thermal conductor at even lower temperatures ^4^.

## Methods

### Finite-element simulations

Time-dependent finite-element simulations of absorption of a radiative beam and heat transfer in solid material were conducted in the COMSOL Multiphysics software Version 6.2 (COMSOL Inc., Burlington, MA). The attenuation of the laser beam was implemented via Beer-Lambert law

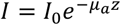

Where *I* and *I*_0_ are the attenuated and initial laser beam power densities, *μ*_*a*_ is the local absorption coefficient of the material that is penetrated by the beam and z is the penetration depth of the laser into the material.

In the finite-elements the equation solved is

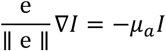

where e is the orientation of the beam.

The absorbed energy at each spatial element is then:

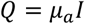

The absorbed energy represents a heat source, which is distributed within the sample via thermal conductivity in the solid materials:

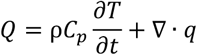

where *ρ* is the density, *C*_*p*_ is the heat capacity at constant pressure and *q* = −*μ*_*a*_∇*T* is the heat flux by conduction.

The geometries of the samples and their mountings were implemented as described in the original publications. Chang et al. used gold EM finder grids covered with a holey carbon film ^44^. These were mounted into the cryostage^2^, a cryostage dedicated to CLEM of EM grids, where the EM grid is mounted to a brass block that is cooled by liquid nitrogen ^1,26^. Since the thermal conductivity of brass (∼110 Wm^-1^K^-1^) and gold (∼310 Wm^-1^K^-1^) are orders of magnitude higher than those of water (<0.6 Wm^-1^K^-1^) or the supporting carbon film (0.5 Wm^-1^K^-1^; ^45^), it was assumed that the brass and the gold adopted the temperature of the liquid nitrogen (−196°C) and are not significantly heated by the laser light. Therefore, one grid square with an edge length of 108 μm was modeled as a layer of water of 500 nm, as given by Chang et al., or 5 μm, which is the approximate height of a mammalian cell. The layer is supported by a carbon film with a thickness of 10 nm (Quantifoil Micro Tools GmbH, Großlöbichau, Germany) and a 2-μm hole in the center (Figure 1a). Alternatively, the sample was supported by a holey gold film ^27^, a holey carbon film coated with 50 nm silver ^21^, 1 or 5 continuous graphene layers ^29^ or a continuous 12-nm SiO_2_ film ^28^. The temperature outside of the square was fixed to −196°C. Above and below the water and carbon are small cavities filled with air, which was modeled as complete insulation, because of the very low thermal conductivity of air (∼0.01 Wm^-1^K^-1^) and because convective effects are largely blocked by the small cavities. Only very minor heat dissipation effects on EM grids have been observed when relatively large convective flows were considered ^23^. Further, the temperature of the air cannot be taken as temperature of the coolant, because it is in contact with the warmer glass windows of the cryostage^2^. Thus, we consider here optimal cooling conditions, since the conduction from the liquid nitrogen over the brass block to the gold EM grid could lead to a small temperature gradient and conduction or convection by the air could lead to minor heating of the whole sample. The cryo-stage developed in ^5^ is dedicated to cryo-arrest during fluorescence live cell imaging and subsequent cryogenic fluorescence microscopy. Here, cells are grown on a standard microscopy cover slide and attached to a heat exchanger made out of chemical vapor deposited diamond that is cooled from the other side by a constant flow of liquid nitrogen. The diamond is mounted in a stainless-steel mount. Initial simulations showed that the stainless-steel mount has negligible effect on the temperature at the irradiated site. Therefore, this sample was modeled as a 10-μm aqueous layer on top of a 150-μm cover slide attached to a beveled diamond disk of 800 μm thickness and 3 mm radius (Figure 1b). The convective heat flux due to the 20°C warm gas at the cover slide and the free parts of the diamond bottom were modeled with a constant heat flux of 5 Wm^-2^K^-1^. The geometries were modeled in 3D using a mesh to discretize the geometry into a finite number of tetrahedral elements. It was made sure to have a high enough number of finite elements in the area of the laser beam and that using a finer mesh did not significantly change the result of the simulation.

SMLM illumination was modeled as described by Chang et al., who illuminated with 0.3 mW of 488-nm laser light and 0.015 mW of 405-nm laser light. Since the absorption of water at 405-nm is approximately 2-fold higher compared to 488-nm laser light, an illumination intensity of 0.33 mW with the water absorption at 488-nm was used. The illumination area was given as 100 μm^2 3^ and was modeled as uniform circular illumination of this area. STED illumination was modeled as a top-hat disk with a radius of 700 nm with a transition zone of 500 nm. The size closely resembles the measured doughnut shaped beam using a 0.95 NA air objective as used in Huebinger et al.^5^ (Supplementary figure 6). The incident power was set to 1 W. This power is achievable in the sample, when commercially available 3-W lasers at 775-nm are used.

Absorption coefficients of water (2.4 m^-1^ at a wavelength of 775-nm; 0.017m^-1^ at 448-nm), thin film amorphous carbon (2.3×10^7^ m^-1^) and the glass cover slide (0.36 m^-1^ at 775-nm) were taken from the literature ^46-50^. Absorption of thin gold film is based on measured values for the real (ε^1^=-2.24) and imaginary (ε^2^=3.98) part of the dielectric function at *λ*=490 nm for a 53-nm gold film ^51^. From this the real (n) and imaginary (*κ*) part of the complex refractive index (ñ) was calculated:

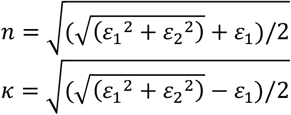

The absorption coefficients (*μ*_*α*_) was then calculated as

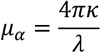

The values for n and *κ* for thin films of silver (n=0.131 and *κ*=2.8), SiO_2_ (n*≈*1.5 and *κ ≈* 0.002) and amorphous carbon (n*≈*2.5 and *κ ≈* 0.375) were taken from the literature ^52-54^. This results in *μ*_*α*_ = 7.2 ∗ 10^7^ m^−1^ and *μ*_*α*_ = 4.7 ∗ 10^7^x m^−1^ and *μ*_*α*_ = 4.5 ∗ 10^4^ m^−1^at *λ* = 488 nm for silver, gold and SiO2, respectively. Reflections were calculated using Fresnels equation for normal incidence:

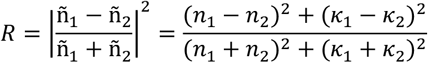

where n_1_=1 and *κ*_1_=0 refer to the values of air. Reflection is incorporated by attenuating the incident laser beam by the reflected fraction.

The transmission (T) of the support material was calculated as:

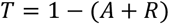

With the absorption (A) in a film of a given thickness (z):

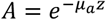

The reported 2.3% of absorption in an atomic layer of graphene ^30^ were directly implemented as a heat source of 100 μm^2^ with 759000 Wm^-2^ in the thermally thin material. Thermal conductivity of the graphene was modelled as asymmetric with 2000 Wm^-1^K^-1^ in plane and 20 Wm^-1^K^-1^ perpendicular to the graphene layer. Free graphene layers can have higher in-plane conductivities ^31^. However, the conductivity is reduced when the layer is in contact with other material and the graphene layers used on EM grids are usually modified graphene layers ^32^ and their exact thermal conductivities were unknown. The absorption coefficient of the chemical vapor deposited diamond at 775-nm (14 m^-1^) was given by the manufacturer (Element Six, London, UK). We used thermal conductivities of thin films of gold (72 Wm^-1^K^-1^) and silver (64 Wm^-1^K^-1^) as calculated by ^21^, based on formulas provided by ^55^, and also used by ^23^.

### Measurement of STED beam size

A sample of 150-nm gold beads embedded in DPX mounting medium (Abberior Instruments GmbH, Göttingen, Germany) was scanned with 20-nm pixel length on a commercial confocal laser-scanning STED microscope (Expert Line; Abberior Instruments GmbH, Göttingen, Germany) equipped with a 775 nm (1.25 W) wavelength STED laser, a vortex phase plate to create the doughnut-shaped laser beam profile, a 40x 0.95NA objective (UPlanApo; Olympus Deutschland GmbH, Hamburg, Germany) and a photomultiplier tube.

### Cell culture

MDCK cells (ATCC No. CCL-34) were obtained from ATCC. The cells were regularly tested for mycoplasma infection using the MycoAlert Mycoplasma detection kit (Lonza, Basel, Switzerland). They were maintained in Dulbecco’s Modified Eagle’s Medium (DMEM) supplemented with 10% fetal bovine serum (FBS), 200 mM L-Glutamine, and 1% nonessential amino acids and cultured at 37 °C with 95% air and 5% CO_2_.

### Determination of absorption coefficient of mammalian cells

MDCK cells were seeded in 2-well silicon culture inserts with a 0.22 cm^2^ area per well (IBIDI GmbH, Gräfelfing, Germany) that were placed into 2-well cell culture chambers on glass cover slides (Sarstedt AG & Co. KG, Nürnbrecht, Germany), as described before ^56,57^. 0.25×10^4^ −1×10^4^ cells were seeded per silicon well, forming a confined monolayer of cells with a clear border after removal of the silicon insert 1-2 days after seeding. The cell culture medium was replaced by 1 mL cell culture medium without phenol red, to have a more transparent medium, and without FCS, to minimize growth factor-induced cell migration ^56,57^.

Transmission images of the border of the monolayer were acquired using an Olympus IX 81 microscope equipped with a 4x 0.16 NA and a 20x 0.7 NA objective and an Orca ER camera. This way the intensity with (*I*) and without (*I*_0_) extinction by the cell monolayer could be measured in the same image (figure 2a). The condenser of the microscope was removed to avoid convergence of the illumination. This way the NA of the objective reflects the collection angle of light (*α*) as *NA* = sin(*α*) for these air objectives (figure 2b). Additionally, wells without cells, but with imaging medium were imaged in order to correct for inhomogeneous illumination.

Afterwards, 10 μL of fluorescein solution was added to the samples and confocal stacks were recorded at a LeicaSP8 microscope equipped with a 60x 1.4 NA oil immersion objective using a white light laser for 488-nm excitation (Supplementary figure 2). From these measurements the height of the monolayer in these samples was measured to be *z* = 5.8 ± 0.3 μm (mean ± sem).

The extinction coefficients (*ε*) were calculated using:

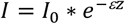

Thus:

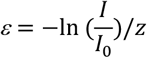

This resulted in *ε*_20*x*_ = 505 ± 29 m^−1^ and *ε*_4*x*_ = 1953 ± 140 m^−1^.

The extinction coefficient is a sum of the absorption coefficient (*μ*_*a*_) and a fraction (x) of the scattering coefficient (*μ*_*s*_) that reflects the light that is deflected strong enough to miss the detector, i.e. miss the objective:

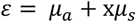

84.6 ± 0.3% and 98 ± 1% of the light scattered by a cell are collected by 4x 0.16NA (*α*=9.2°) and 20x 0.75NA (*α*=48.5) objective^39^ (supplementary figure 3). This results in a solvable set of equations:

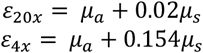

Solving the system of equations:

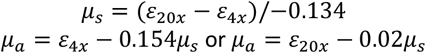

Including standard error propagation, this resulted in *μ*_*s*_ = 10806 ± 1236 *m*^−1^ and *μ*_*a*_ = 289 ± 114 *m*^−1^.

## Acknowledgments

The authors would like to thank Sabrina Seidler and Manuela Grygier for their assistance with the cell culture work as well as Dr. Malte Schmick and Dr. Oliver Hofnagel (all Max-Planck institute of Molecular Physiology) for critical reading of the manuscript.

## Author contribution

J.H. and P.I.H.B. conceived the project. J.H. designed and performed experiments and analysis and wrote the manuscript.

## Data availability statement

All data will be made accessible upon reasonable request to the corresponding author.

## References

1 Sartori, A. et al. Correlative microscopy: Bridging the gap between fluorescence light microscopy and cryo-electron tomography. Journal of Structural Biology 160, 135–145 (2007). 10.1016/j.jsb.2007.07.011

2 Schwartz, C. L., Sarbash, V. I., Ataullakhanov, F. I., Mcintosh, J. R. & Nicastro, D. Cryofluorescence microscopy facilitates correlations between light and cryo-electron microscopy and reduces the rate of photobleaching. Journal of Microscopy 227, 98–109 (2007). 10.1111/j.1365-2818.2007.01794.x

3 Chang, Y.-W. et al. Correlated cryogenic photoactivated localization microscopy and cryo-electron tomography. Nature Methods 11, 737–739 (2014). 10.1038/nmeth.2961

4 Hoffman, D. P. et al. Correlative three-dimensional super-resolution and block-face electron microscopy of whole vitreously frozen cells. Science 367, eaaz5357 (2020). 10.1126/science.aaz5357

5 Huebinger, J. et al. Ultrarapid cryo-arrest of living cells on a microscope enables multiscale imaging of out-of-equilibrium molecular patterns. Science Advances 7, eabk0882 (2021). 10.1126/sciadv.abk0882

6 Dubochet, J. & McDowall, A. Vitrification of pure water for electron microscopy. Journal of Microscopy 124, 3–4 (1981).

7 McDowall, A. W. et al. Electron microscopy of frozen hydrated sections of vitreous ice and vitrified biological samples. Journal of microscopy 131, 1–9 (1983).

8 Macfarlane, D. R. Devitrification in glass-forming aqueous solutions. Cryobiology 23, 230–244 (1986). 10.1016/0011-2240(86)90049-0

9 Mowry, N. J., Krüger, C. R., Drabbels, M. & Lorenz, U. J. Direct measurement of the critical cooling rate for the vitrification of water. Physical Review Research 7, 013095 (2025). 10.1103/PhysRevResearch.7.013095

10 Huebinger, J. et al. Direct Measurement of Water States in Cryopreserved Cells Reveals Tolerance toward Ice Crystallization. Biophysical Journal 110, 840–849 (2016). 10.1016/j.bpj.2015.09.029

11 Voss, J. M., Harder, O. F., Olshin, P. K., Drabbels, M. & Lorenz, U. J. Microsecond melting and revitrification of cryo samples. Structural Dynamics 8 (2021). 10.1063/4.0000129

12 Li, W., Stein, S. C., Gregor, I. & Enderlein, J. Ultra-stable and versatile widefield cryofluorescence microscope for single-molecule localization with sub-nanometer accuracy. Opt. Express 23, 3770 (2015). 10.1364/OE.23.003770

13 Baumeister, W. Cryo-electron tomography: A long journey to the inner space of cells. Cell 185, 2649–2652 (2022). 10.1016/j.cell.2022.06.034

14 Betzig, E. et al. Imaging Intracellular Fluorescent Proteins at Nanometer Resolution. Science 313, 1642–1646 (2006). 10.1126/science.1127344

15 Kaufmann, R. et al. Super-Resolution Microscopy Using Standard Fluorescent Proteins in Intact Cells under Cryo-Conditions. Nano Letters 14, 4171–4175 (2014). 10.1021/nl501870p

16 Liu, B. et al. Three-dimensional super-resolution protein localization correlated with vitrified cellular context. Scientific reports 5, 13017 (2015). 10.1038/srep13017

17 Dahlberg, P. D. et al. Identification of PAmKate as a Red Photoactivatable Fluorescent Protein for Cryogenic Super-Resolution Imaging. J Am Chem Soc 140, 12310–12313 (2018). 10.1021/jacs.8b05960

18 Moser, F. et al. Cryo-SOFI enabling low-dose super-resolution correlative light and electron cryo-microscopy. Proceedings of the National Academy of Sciences 116, 4804–4809 (2019). doi:10.1073/pnas.1810690116

19 Tuijtel, M. W., Koster, A. J., Jakobs, S., Faas, F. G. A. & Sharp, T. H. Correlative cryo super-resolution light and electron microscopy on mammalian cells using fluorescent proteins. Scientific Reports 9, 1369 (2019). 10.1038/s41598-018-37728-8

20 Huebinger, J. Modification of cellular membranes conveys cryoprotection to cells during rapid, non-equilibrium cryopreservation. PLOS ONE 13, e0205520 (2018). 10.1371/journal.pone.0205520

21 Dahlberg, P. D., Perez, D., Hecksel, C. W., Chiu, W. & Moerner, W. E. Metallic support films reduce optical heating in cryogenic correlative light and electron tomography. Journal of Structural Biology 214, 107901 (2022). 10.1016/j.jsb.2022.107901

22 Last, M. G. F., Tuijtel, M. W., Voortman, L. M. & Sharp, T. H. Selecting optimal support grids for super-resolution cryogenic correlated light and electron microscopy. Scientific Reports 13 (2023). 10.1038/s41598-023-35590-x

23 Mojiri, S. et al. Effects of base temperature, immersion medium, and EM grid material on devitrification thresholds in cryogenic optical super-resolution microscopy. Journal of Structural Biology 217, 108231 (2025). 10.1016/j.jsb.2025.108231

24 Hell, S. W. & Wichmann, J. Breaking the diffraction resolution limit by stimulated emission: stimulated-emission-depletion fluorescence microscopy. Optics Letters 19, 780–782 (1994). 10.1364/OL.19.000780

25 Inyushkin, A. V. et al. Thermal conductivity of high purity synthetic single crystal diamonds. Physical Review B 97, 144305 (2018). 10.1103/PhysRevB.97.144305

26 Rigort, A. et al. Micromachining tools and correlative approaches for cellular cryoelectron tomography. Journal of Structural Biology 172, 169–179 (2010). 10.1016/j.jsb.2010.02.011

27 Russo, C. J. & Passmore, L. A. Ultrastable gold substrates for electron cryomicroscopy. Science 346, 1377–1380 (2014). 10.1126/science.1259530

28 Toro-Nahuelpan, M. et al. Tailoring cryo-electron microscopy grids by photomicropatterning for in-cell structural studies. Nature Methods 17, 50–54 (2020). 10.1038/s41592-019-0630-5

29 Russo, C. J. & Passmore, L. A. Controlling protein adsorption on graphene for cryo-EM using low-energy hydrogen plasmas. Nature Methods 11, 649–652 (2014). 10.1038/nmeth.2931

30 Piper, J. R. & Fan, S. Total Absorption in a Graphene Monolayer in the Optical Regime by Critical Coupling with a Photonic Crystal Guided Resonance. ACS Photonics 1, 347–353 (2014). 10.1021/ph400090p

31 Fugallo, G. et al. Thermal Conductivity of Graphene and Graphite: Collective Excitations and Mean Free Paths. Nano Letters 14, 6109–6114 (2014). 10.1021/nl502059f

32 Fujita, J. et al. Epoxidized graphene grid for highly efficient high-resolution cryoEM structural analysis. Scientific Reports 13, 2279 (2023). 10.1038/s41598-023-29396-0

33 Wan, Z. et al. Measuring optical reflectivity of graphene films using compensated Fabry-Perot interferometry. Applied Surface Science 639, 158237 (2023). 10.1016/j.apsusc.2023.158237

34 Mourant, J., Freyer, J. & Johnson, T. Measurements of scattering and absorption in mammalian cell suspensions. Vol. 2679 PW (SPIE, 1996).

35 Cone, M. T. et al. Measuring the absorption coefficient of biological materials using integrating cavity ring-down spectroscopy. Optica 2, 162–168 (2015). 10.1364/OPTICA.2.000162

36 Duyens, L. N. M. The flattering of the absorption spectrum of suspensions, as compared to that of solutions. Biochimica et Biophysica Acta 19, 1–12 (1956). 10.1016/0006-3002(56)90380-8

37 Amesz, J., Duysens, L. N. M. & Brandt, D. C. Methods for measuring and correcting the absorption spectrum of scattering suspensions. Journal of Theoretical Biology 1, 59–74 (1961). 10.1016/0022-5193(61)90026-1

38 Latimer, P. Absolute absorption and scattering spectrophotometry. Archives of Biochemistry and Biophysics 119, 580–581 (1967). 10.1016/0003-9861(67)90493-6

39 Watson, D., Hagen, N., Diver, J., Marchand, P. & Chachisvilis, M. Elastic Light Scattering from Single Cells: Orientational Dynamics in Optical Trap. Biophysical Journal 87, 1298–1306 (2004). 10.1529/biophysj.104.042135

40 Sandell, J. L. & Zhu, T. C. A review of in-vivo optical properties of human tissues and its impact on PDT. Journal of Biophotonics 4, 773–787 (2011). 10.1002/jbio.201100062

41 Dubochet, J., Booy, F. P., Freeman, R., Jones, A. V. & Walter, C. A. Low temperature electron microscopy. Annual review of biophysics and bioengineering 10, 133–149 (1981). 10.1146/annurev.bb.10.060181.001025

42 Franken, L. E., Rosch, R., Laugks, U. & Grünewald, K. Protocol for live-cell fluorescence-guided cryoFIB-milling and electron cryo-tomography of virus-infected cells. STAR Protocols 3, 101696 (2022). 10.1016/j.xpro.2022.101696

43 Berman, R., Foster, E. L., Ziman, J. M. & Simon, F. E. Thermal conduction in artificial sapphire crystals at low temperatures I. Nearly perfect crystals. Proceedings of the Royal Society of London. Series A. Mathematical and Physical Sciences 231, 130–144 (1955). 10.1098/rspa.1955.0161

44 Ermantraut, E., Wohlfart, K. & Tichelaar, W. Perforated support foils with pre-defined hole size, shape and arrangement. Ultramicroscopy 74, 75–81 (1998). 10.1016/S0304-3991(98)00025-4

45 Bullen, A. J., O’Hara, K. E., Cahill, D. G., Monteiro, O. & von Keudell, A. Thermal conductivity of amorphous carbon thin films. Journal of Applied Physics 88, 6317–6320 (2000). 10.1063/1.1314301

46 Palmer, K. F. & Williams, D. Optical properties of water in the near infrared*. J. Opt. Soc. Am. 64, 1107–1110 (1974). 10.1364/JOSA.64.001107

47 Hass, M. & Davisson, J. W. Absorption coefficient of pure water at 488 and 541.5 nm by adiabatic laser calorimetry*. J. Opt. Soc. Am. 67, 622–624 (1977). 10.1364/JOSA.67.000622

48 Bassarab, V. V., Shalygin, V. A., Shakhmin, A. A., Sokolov, V. S. & Kropotov, G. I. Spectroscopy of a borosilicate crown glass in the wavelength range of 0.2 µm–15 cm. Journal of Optics 25, 065401 (2023). 10.1088/2040-8986/accaf9

49 Agar, A. W. The measurement of the thickness of thin carbon films. British Journal of Applied Physics 8, 35 (1957). 10.1088/0508-3443/8/1/310

50 Moretz, R. C., Johnson, H. M. & Parsons, D. F. Thickness estimation of carbon films by electron microscopy of transverse sections and optical density measurements. Journal of Applied Physics 39, 5421–5424 (1968). 10.1063/1.1655992

51 Yakubovsky, D. I., Arsenin, A. V., Stebunov, Y. V., Fedyanin, D. Y. & Volkov, V. S. Optical constants and structural properties of thin gold films. Opt. Express 25, 25574–25587 (2017). 10.1364/OE.25.025574

52 Lynch, D. W. & Hunter, W. R. in Handbook of Optical Constants of Solids (ed Edward D. Palik) 275–367 (Academic Press, 1985).

53 Rodríguez-de Marcos, L. V., Larruquert, J. I., Méndez, J. A. & Aznárez, J. A. Self-consistent optical constants of SiO2 and Ta2O5 films. Opt. Mater. Express 6, 3622–3637 (2016). 10.1364/OME.6.003622

54 Skowronski, L., Chodun, R., Trzcinski, M. & Zdunek, K. Optical Properties of Amorphous Carbon Thin Films Fabricated Using a High-Energy-Impulse Magnetron-Sputtering Technique. Materials (Basel) 16 (2023). 10.3390/ma16217049

55 Kumar, S. & Vradis, G. C. Thermal Conductivity of Thin Metallic Films. Journal of Heat Transfer 116, 28–34 (1994). 10.1115/1.2910879

56 Brüggemann, Y., Karajannis, L. S., Stanoev, A., Stallaert, W. & Bastiaens, P. I. H. Growth factor-dependent ErbB vesicular dynamics couple receptor signaling to spatially and functionally distinct Erk pools. Science Signaling 14, eabd9943 (2021). 10.1126/scisignal.abd9943

57 Joshi, M. S. et al. The EGFR phosphatase RPTPγ is a redox-regulated suppressor of promigratory signaling. The EMBO Journal 42, e111806 (2023). 10.15252/embj.2022111806

