## Supplementary Figures for "Determination of factors that allow cryogenic nanoscopy with high power illumination without devitrification"

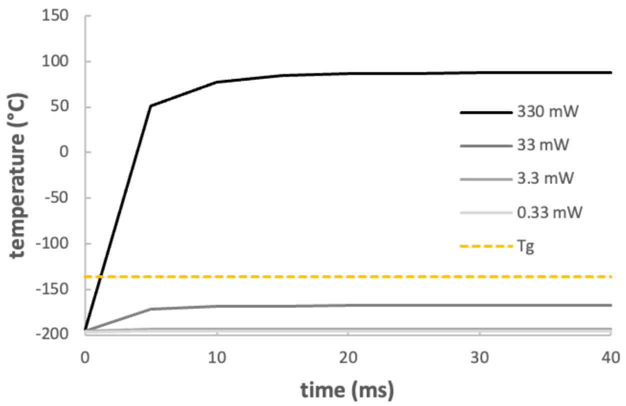


**Supplementary figure 1: Heating of water sample by SMLM on a $SiO_2$ support**
**Temperature over time in the central point of the water layer from start of the 488-nm laser irradiation at a 100-µm$^2$ (3.3x10$^2$ – 3.3x10$^5$ W/cm$^2$) spot in the center in the complete EM grid configuration using a 12-nm $SiO_2$ support and different laser intensities (gray lines); Tg: glass transition temperature of water (dashed yellow line).**

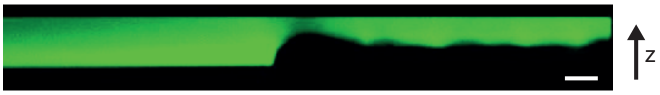


**Supplementary figure 2: Cell thickness measurement by confocal laser scanning microscopy**
**Representative image of a thickness measurement of a monolayer of MDCK cells. After addition of fluorescein solution, confocal stacks of the edge of confined monolayers of MDCK cells used for extinction measurements (figure 3a) were recorded. Shown is a reconstruction of an xz-plane that was used to measure the thickness of the cell layer along the z-axis. Scale bar: 10 µm**

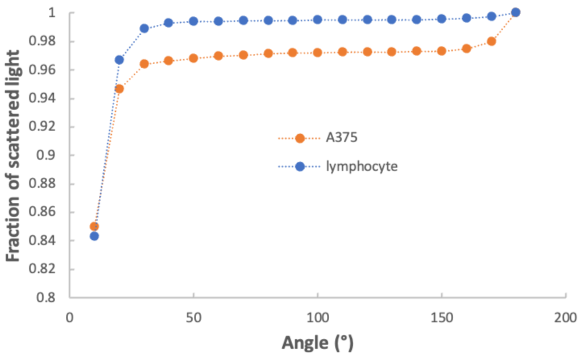


**Supplementary figure 3: Angle-dependent scattering of mammalian cells**

**Scattering intensities over scattering angle were extracted from Watson et al., 2004, Biophysical Journal, Figures 6 and 12. The missing last 20° (A375) or 30° (lymphocyte) values were extrapolated from the previous 30° using exponential functions with $R^2$=0.999 and 0.97, respectively. Areas under the curves were calculated every 10°.**

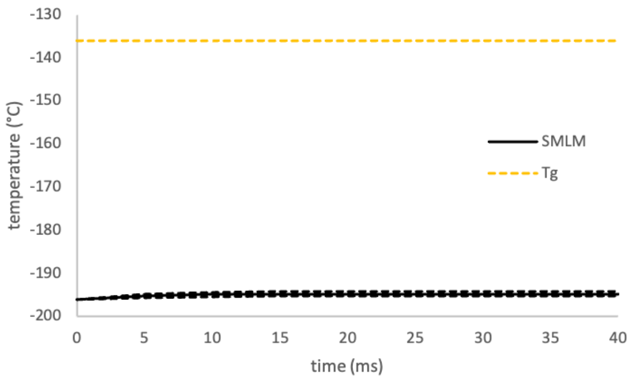


**Supplementary figure 4: Heating of a cellular sample by SMLM on an EM grid**
**Temperature over time in the central point of the cell layer (absorption coefficient 289 $\pm$ 114 m$^{-1}$) from start of the 488-nm laser irradiation at a 100-$\mu$m$^2$ spot in the center of the complete EM grid configuration omitting the support film (black lines; mean $\pm$ sem); Tg: glass transition temperature of water (dashed yellow line).**

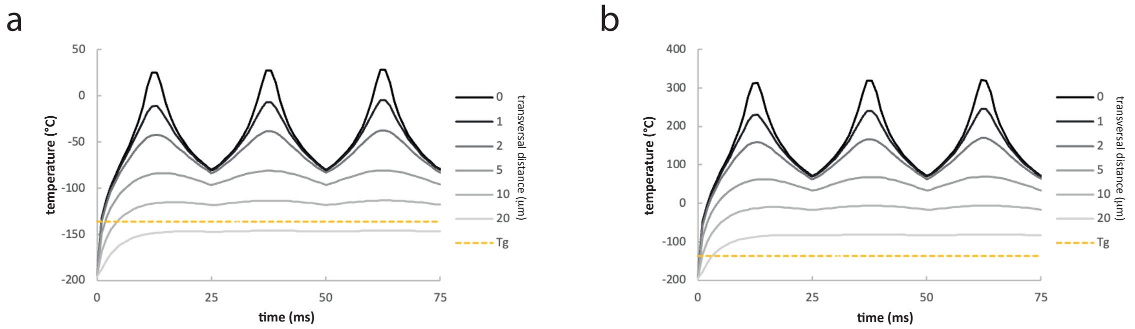


**Supplementary figure 5: Temperature course during scanning of cells on an EM grid**

**Temperature curse during scanning of 3 10-µm lines by the 1-W STED laser at -40 nm, 0 nm and +40 nm in transversal direction relative to the measurement point over a cellular sample with the minimum determined absorption coefficient of 175 $m^{-1}$ and the maximum determined absorption coefficient of 403 $m^{-1}$ (b). The temperature is depicted at the center of the 2$^{nd}$ scanning line and at indicated transversal distances to the measurement point (gray lines). Tg: glass transition temperature of water (yellow dashed line)**

a b

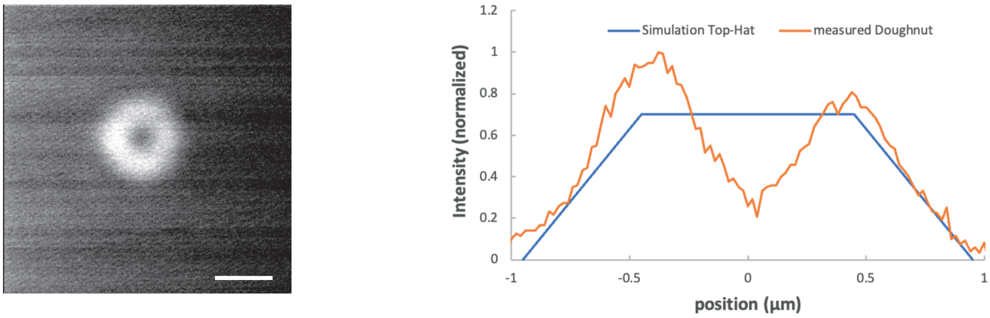


**Supplementary figure 6: STED laser beam size**

**a) The doughnut-shaped illumination profile of a was measured by reflection from a 150-nm gold particle using a 40x 0.95 NA objective. Scale bar: 1 µm b) A background-corrected line profile through the doughnut-shape in a) is compared to the top-hat shape that is used for the simulation. Both profiles have been normalized to an area under the curve of 1.**

**Supplementary Movie 1: Heating of an aqueous sample on an EM grid**
**Shown is the temperature following irradiation of a circular region of 100 µm$^2$ in the center of the gold EM grid mesh with a holey carbon support film by 0.33 mW of 488-nm light over 40 ms as simulated by finite-element simulation. Temperature is color-coded as indicated on the right.**

**Supplementary Movie 2: Heating of a cell sample on an EM grid by a scanning STED laser**
**Shown is the irradiation intensity (left, color-code in W/m$^2$) and the temperature (right, color-code in °C) following scanning of a 1-W STED laser focused to a radius of 0.7 µm over three 10-µm lines with a scanning speed to acquire 40-nm pixel length with a pixel dwell time of 100 µs as simulated by finite-element simulation. The temperature was simulated by absorption of the laser light by the cells with an absorption coefficient of 289 m$^{-1}$ without considering absorption by a support film. The lines were scanned in the center of a gold EM grid mesh (see figure1).**
